# Multiple Postsynaptic Protein Levels in Adult Superior Colliculus Are Unaffected by Dark Rearing from Birth

**DOI:** 10.1101/2022.10.06.511220

**Authors:** Parag S. Juvale, David B. Mudd, Nitheyaa Shree, Sarah L. Pallas

**Affiliations:** Department of Biology, University of Massachusetts Amherst, Amherst, MA 01003, USA; Neuroscience Institute, Georgia State University, Atlanta, GA 30303, USA

**Author notes:** Correspondence: Sarah L. Pallas Department of Biology, University of Massachusetts-Amherst Amherst, MA 01003 USA. Office of Technology Transfer, Emory University, Atlanta, GA. Graduate Program in Biochemistry, University of Bristol, UK.

**Keywords:** Visual deprivation, superior colliculus, GABA, adult plasticity, visual refinement, GABA_A_ receptor, gephyrin, chloride co-transporter

## Abstract

Visual deprivation by dark rearing in kittens and monkeys delays visual pathway development and prolongs the critical period. In contrast, receptive fields (RFs) in superior colliculus (SC) of Syrian hamsters *(Mesocricetus auratus)* refine normally with spontaneous activity alone, requiring only brief juvenile visual experience to maintain refined RFs in adulthood (Carrasco et al., 2005). Extending dark rearing past puberty leads to lower GAD and GABA levels due to reduced BDNF-TrkB signaling, resulting in RF re-enlargement (Carrasco et al., 2011; Mudd et al., 2019). Previous studies in kittens and monkeys have reported that dark rearing is associated with changes in both GABA ligand and GABA_A_ receptor levels. Given the reduced GABA levels in SC of dark reared adult hamsters, we asked if dark rearing also causes changes in GABA_A_ receptor levels. We examined expression of GABA_A_ receptor subunits, their anchoring protein gephyrin, and the cation-chloride co-transporters KCC2 and NKCC1 in dark reared hamsters. Surprisingly, we found that dark rearing from birth until puberty had no effect on the levels of any of these postsynaptic elements, revealing a new form of maladaptive, presynaptic only inhibitory plasticity in which, rather than extending the critical period as seen in kittens and monkeys, hamster receptive fields refine normally and then lose refinement in adulthood. These results suggest that attempts to increase plasticity in adulthood for rehabilitation or recovery from injury should consider the possibility of unintended negative consequences. In addition, our results demonstrate the interesting finding that changes in neurotransmitter levels are not necessarily coordinated with changes in postsynaptic components.

## Introduction

During brain development, synaptic connections are elaborated and then refined to a mature state under the influence of neural activity. Activity due to sensory experience during an early “critical period” is important for shaping some aspects of neural circuit development. In the visual pathway, although spontaneous retinal activity is important for initial axon pruning (Kutsarova et al., 2017), it has long been thought that early visual experience is essential to attain refined connectivity patterns during development (Wiesel and Hubel, 1965; Maffei and Galli-Resta, 1990; Meister et al., 1991; Wong et al., 1993; Katz and Shatz, 1996; Firth et al., 2005). Once critical periods have closed, plasticity is often limited or even prevented, thus protecting refined circuits from destabilization (Hubel and Wiesel, 1970; Takesian and Hensch, 2013; Hübener and Bonhoeffer, 2014; Pallas, 2017; Hensch and Quinlan, 2018; Reh et al., 2020; Ribic, 2020; Mitchell and Maurer, 2022).

In some mammals, dark rearing is reported to delay or prevent refinement, prolonging critical period plasticity (Cynader and Mitchell, 1980; Mower et al., 1985; Mower, 1991; Lee and Nedivi, 2002; Nakadate et al., 2012). In apparent contradiction to this common view, we have reported that spontaneous activity is sufficient for refinement of receptive fields in both visual cortex (V1) and superior colliculus (SC) of dark reared (DR) Syrian hamsters *(Mesocricetus auratus)*, and early light exposure for 3-7 days is necessary only to maintain the refinement into adulthood (Carrasco et al., 2005; Carrasco and Pallas, 2006; Balmer and Pallas, 2015; Mudd et al., 2019). Syrian hamster pups spend the first 3-4 weeks after birth underground in the wild (Adler, 1948; Nowosielski-Slepowron and Park, 1987), so it would not be beneficial to have the development of their visual function depend on light exposure. If early visual experience continues to be unavailable, GABAergic lateral inhibition in SC and V1 declines and RFs expand, but not until approximately two months of age (∼puberty) (Carrasco et al., 2005; Balmer and Pallas, 2015; Mudd et al., 2019). Pharmacological activation of TrkB receptors can mimic the effects of early light exposure in DR hamsters, resulting in long-term maintenance of refined receptive fields and visual acuity (Mudd et al., 2019), perhaps through promoting GABA synthesis (Zhang et al., 2018). Thus, hamsters, contrary to what has been found in cats and monkeys, need visual experience to maintain refined receptive fields in adulthood, but not to refine them during development (**Figure 1**).

**Figure 1:** Differences in the visual refinement of hamster visual pathways depending on exposure to light during the critical period of heightened neural plasticity. Animals exposed to light during postnatal development gradually improve RF refinement in SC and V1 and maintain it throughout life (as indicated by the blue line). Animals that do not experience postnatal light also show RF refinement by P60, but the refinement progressively declines in adulthood (as indicated by the orange line).

GABA_A_ receptors are pentameric, ionotrophic receptors consisting of five subunits grouped around a central chloride ion pore. The functional characteristics of the receptor largely depend upon the subunit composition (Sigel et al., 1990) and organization (Minier and Sigel, 2004). Of the many subunit arrangements, alpha1 and alpha2 subunits have been linked to synaptic localization of GABA_A_ receptors. However, these two subunit types have different kinetics and are expressed at different points in development. At birth, receptors containing the alpha2 subunit are widely expressed throughout the brain, whereas alpha1 expression is initially low in major areas of the brain like the neocortex, the hippocampus, and the cerebellum (Laurie et al., 1992; Fritschy et al., 1994; Dunning et al., 1999; Chen et al., 2001). During the first several postnatal weeks, alpha1 expression increases, coinciding with a reduction in alpha2 expression (Fritschy et al., 1994). This alpha2 to alpha1 switch in subunit expression underlies a developmental decrease in inhibitory postsynaptic current (ipsc) decay time and an increase in ipsc amplitude (Fritschy et al., 1994; Okada et al., 2000; Yu et al., 2006).

Our lab demonstrated previously that the expansion of RFs in SC of adult, DR hamsters is associated with a loss of GABA immunoreactivity (Carrasco et al., 2011). Iontophoretic application of GABA_A_ agonists *in vivo* restored RFs to a normal adult size. In addition to a loss of GABA-immunoreactivity, GABA_A_ agonists and antagonists were less effective in SC and V1 neurons of DR hamsters than in normally reared (NR) hamsters (Carrasco et al., 2011).

The incomplete development of RF properties in V1 of visually deprived cats has been associated with a failure in developmental maturation of NMDA and GABA_A_ receptors (Carmignoto and Vicini, 1992; Chen et al., 2000; Chen et al., 2001; Erisir and Harris, 2003). Although a failure to maintain refined RFs is a different phenomenon than a failure to refine them during a critical period, the mechanism(s) could be similar or convergent. We thus tested the hypothesis that maintenance of refined RFs in adulthood depends on the stability of mature receptors and other postsynaptic signaling components. This hypothesis predicts that the detrimental, post-critical period receptive field plasticity observed in DR adult hamsters results from a deprivation-induced failure to maintain these postsynaptic proteins in their mature state. Contrary to this hypothesis, we find, using Western blot assays of synaptosomes, that the quantity, subunit composition, and localization of GABA_A_ receptors in SC of adult dark reared hamsters with re-expanded RFs resemble those seen in normally reared subjects. Furthermore, levels of the synaptic scaffolding proteins gephyrin and PSD-95 are normal, as are the adult expression levels of cation-chloride co-transporters (KCC2/NKCC) in DR subjects. These findings suggest that, although a change in effectiveness of GABA_A_ receptors was reported previously using pharmacological agents (Carrasco et al., 2011; Balmer and Pallas, 2015), the loss of RF refinement in adulthood is mediated primarily by reductions in GABA expression in the presynaptic terminals rather than by significant postsynaptic alterations. This result is at odds with the common view that presynaptic changes in the ligand must occur together with corresponding postsynaptic changes in receptor levels (Fisher-Lavie and Ziv, 2013; Sudhof, 2018 ; Sanderson et al., 2020). Taken together, our results rule out several hypotheses about the mechanistic basis of refined RF maintenance throughout adulthood and provide insights into regulation of critical period plasticity that could help to understand the regulation of GABAergic signaling at the synaptic level. Similar research in diurnal animals that have photopic vision as do humans could help to provide insight on treatment and therapeutic modalities in adults facing issues with plasticity in adulthood.

## Materials and Methods

### Subjects

A total of 42 adult Syrian hamsters (Cricetidae: *Mesocricetus auratus*) (aged P90-P100) of both sexes were bred within our animal facility and used as subjects in this study. Syrian hamsters are an altricial rodent species that is ideal for studying the developing visual system due to their robust and well-characterized visual responses, short gestation time, and large litters (Chalupa, 1981; Huck et al., 1988; Pratt and Lisk, 1989; Razak et al., 2003; Carrasco et al., 2005). Sexual maturity in this species occurs between postnatal days (P)56 and P60 (Diamond and Yanagimachi, 1970; Fitzgerald and Zucker, 1976). Breeding females were singly housed. Male breeders were introduced and supervised until intromission was observed, after which they were removed. Weanlings and adult males were housed in single sex social groups of up to 5 adults per cage and adult females were housed with female siblings or individually in standard rodent cages. Running wheels were not provided because they have been shown to alter the timing of visual cortical plasticity (Baroncelli et al., 2010; Tognini et al., 2012; Kalogeraki et al., 2014) but a variety of other enrichment items were available. All subjects were provided with *ad libitum* access to rodent chow and water.

### Treatment groups

Animals used in this study were bred in-house to control sensory experiences throughout development. Dams of DR subjects were transferred into total darkness 1-3 days before parturition. An antechamber and light-impenetrable black curtain separated the dark housing room from the hallway, ensuring that any accidental openings of the hallway doors did not expose the animals to light. Dark reared animals were housed inside light-tight stackable cages with a standard HVAC filtration system consistent with the other animal rooms in the facility. During general animal husbandry purposes, the hamsters were exposed to dim red light at a wavelength not visible to Syrian hamsters (Huhman and Albers, 1994). NR hamsters were maintained in a standard 12/12 light cycle room from before birth into adulthood.

### Western blotting

Animals were euthanized with Euthasol at >150 mg/kg IP prior to tissue collection. Brains were immediately extracted and frozen in 2-methylbutane on dry ice, then stored at -80°C or immediately dissected for preparation of lysates. In order to analyze differences in protein levels between NR and DR hamsters, we used immunoblotting (Western blots). Western blots can detect protein levels at a 1-3 ng resolution (Coorssen et al., 2002; Ghosh et al., 2014), allowing high resolution quantification of proteins. Note that normal levels of synaptic GABA concentration have been estimated to be between 1.5-3 mM (Coorssen et al., 2002; Tretter et al., 2008; Ghosh et al., 2014). GAPDH or β-actin were used as loading controls to normalize for any differences in the amount of lysate pipetted into each gel lane.

Protein extraction was done as described by Shi et al. (1997) with a few modifications. Briefly, individual left and right SC brain areas were excised and homogenized in a lysis buffer (10 mM phosphate buffer, pH 7.0, 5 mM EGTA, 5 mM EDTA, 1 mM DTT) containing Halt protease inhibitor (ThermoFisher Scientific). The lysate was centrifuged at 13,000 rpm at 4°C for 10 min, and the supernatant was saved for the analysis of cytosolic proteins (cytosolic fraction). The resulting pellet was resuspended in 2 mM HEPES buffer, pH 7.2. It was then centrifuged at 70,000 rpm at 4°C for 30 min. The pellet thus obtained was resuspended in 0.5 mM HEPES, pH 7.3, containing 0.32 M sucrose and centrifuged at 2,000 rpm for 8 min. Synaptosomes are present in the supernatant with this method. The success of the synaptosome isolation protocol was confirmed by assessing the presence of Histone H3, which should only be present in the cytosolic fractions and not in the synaptosome fractions (**Figure 2**). Synaptic proteins were then quantified using the Pierce BCA Protein Assay Kit (ThermoFisher Scientific) mixed with 2X sample buffer and heated for 15 min at 60 °C. Twenty μg of the synaptosome proteins were loaded per well in pre-cast Bio-Rad gels and electrophoresis was carried out at 110 V for 90 min in a Bio-Rad electrophoresis tank. Proteins were then transferred onto nitrocellulose membranes at 70 V for 75 min, blocked in BSA for 1 h, and probed with primary antibodies overnight. Primary antibodies used in this study included: rabbit anti-GABA_A_Rα1 (1:1000, Cat#: AGA-001, Alomone Labs); rabbit anti-GABA_A_Rα2 (1:1000, Cat#: ab72445, Abcam); rabbit anti-GABA_A_Rα5 (1:1000, Cat#: ab10098, Abcam); rabbit anti-Gephyrin (1:1000, Cat#: ab32206, Abcam); mouse anti-PSD-95 (1:1000, Cat#: ab2723, Abcam); mouse anti-KCC2 (1:1000, Cat#: 75-013, NeuroMab); rabbit anti-NKCC1 (1:1000, Cat#: ab59791, Abcam); mouse anti-β-actin (1:1000, Cat#: A2228, Sigma-Aldrich) and mouse anti-GAPDH (1:1000, Cat#: 600-GAPDH, PhosphoSolutions). Protein bands were labeled using either appropriate fluorescent secondaries or appropriate HRP-conjugated secondary antibodies, then imaged on an Odyssey CLx fluorescent imaging system (Li-Cor) or developed with enhanced chemiluminescent (ECL) substrate in a Bio-Rad ChemiDoc Imager. All of the proteins studied here were analyzed and quantified as a ratio of the optical density of the protein of interest compared to the density of the loading control (either GAPDH or β-actin). Note that GAPDH is found in the pre-and post-synaptic sites (Frederikse et al., 2016) along with β-actin, and thus makes an effective control protein to measure relative densities.

**Figure 2.** Histone H3 expression in the synaptosomal and cytosolic lysates. Western blot showing histone H3 (15 kDa) bands in the synaptosomal and cytosolic fractions of SC. GAPDH was used as a loading control. The presence of histone H3 in the cytosolic fraction and its absence in the synaptosomal fraction shows an effective synaptosomal separation occurred in these experiments.

### Statistical Analysis

A Student’s *t*-test was used to compare parametric data with equal variance between treatment groups and a normally distributed control data set. In the case of non-parametric data (data that were not normally distributed and/or exhibited unequal variance), a Mann-Whitney Rank Sum (U) test was employed. Descriptive statistics for these analyses are provided as mean ± standard error of the mean (SEM) in the text. Whiskers represent the 5th (lower) and 95th (upper) percentage of the data.

## Results

The failure to maintain RF refinement in adult DR animals involves deficits in overall GABA expression and GABA_A_ receptor function (Carrasco et al., 2011), leading to a loss of lateral inhibition and thus expansion of RFs after P60 (Carrasco et al., 2005; Balmer and Pallas, 2015; Mudd et al., 2019). Using adult hamsters (postnatal day (P)90-P100), we explored several possible ways that dark rearing during a critical period for RF refinement could affect levels of GABA_A_ receptors and other postsynaptic proteins associated with inhibitory synaptic function in adult SC. We used Western blotting on synaptosomes in normal and dark reared animals to study postsynaptic proteins that might regulate synaptic plasticity.

### Dark rearing does not affect the subunit composition of GABA_A_ receptors in adult SC

Deprivation-induced decreases in both GABA_A_ and NMDA receptor levels in cat visual cortex have been reported previously and were proposed to be involved in functional deficits (Carmignoto and Vicini, 1992; Chen et al., 2000; Chen et al., 2001). In DR hamsters, GAD and GABA immunoreactivity declines (Carrasco et al., 2011; Otfinoski & Pallas, in prep.) and GABA_A_ receptors are less efficient when tested pharmacologically (Carrasco et al., 2011), thus we expected to see changes in levels of postsynaptic GABA_A_ proteins. However, in our previous study using Western blots on synaptosomes, no significant changes in the level of the GABA_A_ receptor alpha1 subunit were observed (Mudd et al., 2019). These results raised the question of whether subunit composition of GABA_A_Rs might be altered by dark rearing in a way that reduced their effectiveness without affecting alpha1 levels.

We reasoned that the developmental alpha2 to alpha1 switch, if reversed in adulthood, could underlie the reduction in GABA_A_ receptor function that was previously observed in studies of RF enlargement in adult SC (Carrasco et al., 2011). We explored this possibility by examining the expression of each subunit in synaptosomes of SC in normally reared and visually deprived adults. Hamsters in the experimental group were dark reared from before birth. We used Western blotting for a high resolution, quantitative assay of synaptic membrane-bound alpha1 and alpha2 GABA_A_ receptor expression in the synaptosome fractions obtained from adult SC. We found that there were no significant differences in either the overall levels of alpha2 protein, observed as a ratio of alpha2 to GAPDH (NR: 1.21 ± 0.052, n=8; DR: 1.33 ± 0.185, n=8; U=26, n=8, p=0.574; Mann-Whitney Rank Sum Test) (**Figure 3A**), or in the ratio of alpha1/alpha2 expression, observed as a ratio of the normalized alpha 1 density (alpha 1/GAPDH ratio) to normalized alpha 2 density (alpha 2/GAPDH ratio) in the SC of adult DR (1.651 ± 0.277, n=5) compared to adult NR hamsters (1.19± 0.084, n=4) (T(7)= -1.436, n=4, p=0.194 Student’s t-test) (**Figure 3B**). This was a surprising result considering our previous finding that dark rearing reduces the response to pharmacological application of GABA agonists and antagonists (Carrasco et al., 2011). These findings argue against a reversal of the normal developmental transition from alpha2 to alpha1-dominant expression as a cause of the deprivation-induced RF enlargement in adult SC, and they support the interpretation that visual experience is not needed to maintain mature GABA_A_ receptor alpha1/alpha2 subunit composition.

**Figure 3.** GABA_A_α2 receptor subunit levels in SC are not affected by early dark rearing. **(A) Image**: Individual Western blots of normally reared (NR) and dark-reared (DR) treatment groups generated using 20 µg of SC protein per lane. GAPDH was used as a loading control. **Plot:** Boxplot showing the normalized GABA_A_Rα2 expression level in normally reared vs. dark reared hamsters. **(B) Image:** Individual Western blots of SC tissue from NR and DR animals comparing GABA_A_Rα1 and GABA_A_Rα2 expression with corresponding GAPDH expression. **Plot:** Boxplot showing the ratio of normalized values of GABA_A_Rα1/GAPDH to the normalized values of GABA_A_Rα2/GAPDH. Boxes in each individual boxplot show the median and 25^th^ and 75^th^ percentiles of the data (whiskers show 5% and 95% levels). Individual data points obtained from each animal within a group are shown as dots. For Western blots in this and the following figures, lanes presented together are from the same gel, and each measured protein was normalized against GAPDH (unless stated otherwise) as a loading control. Taken together, these results reveal that the levels and ratio of synaptic GABA_A_Rα2 receptor subunits and their ratio with GABA_A_Rα1subunits are similar in normal and dark reared adult SC.

GABA_A_ receptors can also be expressed extrasynaptically, where they can be activated by GABA derived from synaptic spillover or non-neuronal sources. This low concentration GABA source generates “tonic” inhibition (Farrant and Nusser, 2005). Alpha5 subunit-containing receptors are primarily expressed extrasynaptically and have been implicated in regulating the induction of synaptic plasticity for LTP in hippocampus (Saab et al., 2010; Zurek et al., 2012; Zurek et al., 2014; Jacob, 2019). However, alpha5 GABA_A_Rs can relocate to the synapse and colocalize with gephyrin (Brady and Jacob, 2015). To investigate the possible role of synaptic alpha5 levels in adult RF maintenance we quantified and compared the alpha5/GAPDH ratios between NR (0.544 ± 0.0520, n=8) and DR (0.471 ± 0.0935, n=8) adult hamsters (**Figure 4A**) using Western blotting. We found no significant differences in alpha5 protein levels between groups (U (20) = 0.308, p = 0.878, Mann-Whitney Rank Sum test). We compared the ratio of alpha1/alpha5 between adult NR (1.027 ± 0.0815, n=10) and DR (0.995 ± 0.0926, n =9) hamsters and found no differences between these groups (U (18) =35, p=0.438, Mann-Whitney Rank Sum test) (**Figure 4B**). Because we were only studying proteins from synaptosome preparations (i.e., GABA_A_ receptor subunits localized in the synapses), these results suggest that the localization of alpha1 and alpha5 subunit-containing GABA_A_ receptors in SC is not being altered by early visual experience.

**Figure 4.** GABA_A_α5 receptor subunit levels in SC are not affected by early dark rearing. A and B Images: Representative Western blots of NR and DR treatment groups as in Figure 3. All lanes presented together are from the same gel(s), and each gel was run with GAPDH as a loading control. **(A) Plot:** Boxplot showing normalized GABA_A_Rα5 expression levels compared between normal and dark reared hamsters. **(B) Plot:** Boxplot showing the ratio of normalized (against GAPDH levels) GABA_A_Rα1: GABA_A_Rα5 expression ratios. Boxes in each individual boxplot show the median and 25^th^ and 75^th^ percentiles of the data, with whiskers at 5 and 95%. Individual data points in each group are shown as dots. These results show that the levels of synaptic GABA_A_α5 receptor subunits and their ratio with GABA_A_Rα1 subunits are similar in normal and dark reared adult SC.

### Dark rearing does not affect the normal location of GABA_A_ receptors in adult SC

The regulation of GABA_A_ receptors at the synapse is pivotal for maintaining correct levels of inhibitory synaptic transmission (Jacob et al., 2008). Impaired trafficking of GABA_A_ receptors into and out of synaptic membranes could affect their synaptic localization in SC and thus their overall response to presynaptically released GABA. GABA_A_ receptor trafficking is partially regulated by endocytosis – the controlled removal of receptors from the postsynaptic membrane into the cytoplasm (see Lévi and Triller, 2006, for review). Endocytosed receptors are subsequently reinserted into the postsynaptic membrane or undergo lysosomal degradation (Kittler et al., 2000). We reasoned that if internalization was dysregulated, either by decreased receptor reinsertion or increased receptor degradation, it could negatively impact the efficacy of GABA_A_ receptors at the synapse. We examined this possibility by comparing the ratio of postsynaptic membrane-bound to cytosolic alpha1 subunit-containing receptors between our treatment groups. No differences were observed in the postsynaptic membrane/cytosol ratio of alpha1 expressing receptors between DR (0.768 ± 0.044, n=7) and NR (0.778 ± 0.1, n=6) adult hamsters (U(11)= 20.00, p=0.945, Mann-Whitney Rank Sum test) (**Figure 5A**). These results indicate that the overall trafficking of alpha1 subunit-expressing synaptic GABA_A_ receptors is not affected by dark rearing.

**Figure 5.** Internalization of GABA_A_ receptors in SC is not affected by early dark rearing. **(A)** Adult levels of the cytosolic vs. the synaptic membrane-attached ratio of the synaptically-targeted GABA_A_Rα1 and **(B)** the synaptically-targeted GABA_A_Rα5 subunits were not affected by early light deprivation. **Images:** Representative Western blots represent bands of cytoplasmic and membrane bound receptor subunit proteins, each from the same animal, measured as a ratio against GAPDH and compared between NR and DR groups. **Plots:** Boxplot showing the ratio of normalized (against GAPDH) values of cytosol: membrane ratios of each subunit. Boxes in each individual boxplot show the median and 25^th^ and 75^th^ percentile of the data, with whiskers indicating 5 and 95%. Individual data points obtained in each group are shown as dots. These results show that the internalization of synaptic GABA_A_Rα1 and GABA_A_Rα5 subunits is similar in normal and dark reared adult SC.

We also examined the possibility that extrasynaptic alpha5 receptor internalization may be dysregulated and thus responsible for changes in tonic GABA_A_ inhibition (Davenport et al., 2021). There were no significant differences in cytosolic membrane localization of alpha5 subunits between adult DR (1.320 ± 0.198, n=8) and NR groups, however (1.753 ± 0.449, n=6) (U(12)=19.00, p=0.573, Mann-Whitney Rank Sum test) (**Figure 5B**). These results indicate that the internalization of extrasynaptic alpha5 subunit-expressing GABA_A_ receptors is not responsible for the decreased efficacy of GABA_A_ receptors observed in RFs that fail to maintain refinement in adulthood after dark rearing.

### Inhibitory and excitatory scaffolding proteins in SC are not affected by dark rearing

One factor influencing the accumulation and retention of GABA_A_ receptors at postsynaptic sites is the membrane scaffolding protein gephyrin (Kneussel et al., 1999; Sun et al., 2004; Jacob et al., 2005; Tretter et al., 2008). Decreased expression of gephyrin results in less clustering (Essrich et al., 1998) and more mobility of GABA_A_ receptors at the synapse (Jacob et al., 2005). We surmised that decreased gephyrin expression could be responsible for the weaker GABA_A_ receptor signaling observed in neurons with RFs that failed to maintain refinement in adulthood. We quantified and compared postsynaptic membrane-bound gephyrin expression between DR and NR adults using Western blotting. Gephyrin levels in DR adults (0.786 ± 0.124, n=17) were similar to those in NR adults (0.736 ± 0.158, n=16) (U(111)=-0.247, p=0.377, Mann-Whitney Rank Sum test) (**Figure 6A**). This indicates that maintenance of adult gephyrin expression levels is not affected by dark rearing and suggests that if GABA_A_ receptor accumulation and trafficking is being affected, then it is occurring independently of gephyrin levels.

**Figure 6.** Gephyrin and PSD-95 expression in SC are not affected by dark rearing. **Images: (A)** Gephyrin and **(B)** PSD-95 expression was similar between adult NR and DR groups (upper panels). **Plots:** Boxplot showing the ratio of normalized values of gephyrin vs. GAPDH (A) or β-actin (B) . Boxes in each individual boxplot show the median and 25^th^ and 75^th^ percentiles of the data, with whiskers at 5 and 95%. Individual data points obtained in each group are shown as dots. These results show that the levels and ratio of the scaffold proteins are similar in normal and dark reared adult SC.

PSD-95 is the primary glutamate (AMPA and NMDA) receptor scaffolding protein in CNS neurons (Chen et al., 2015), and it functions like gephyrin does for GABA_A_ receptors. Although it does not directly impact GABA_A_ receptor function, PSD-95 has an influence on visual circuit plasticity. For example, mice lacking PSD-95 have lifelong ocular dominance plasticity in primary visual cortex that results from an increase in the overall proportion of silent synapses, despite having normal inhibitory tone (Funahashi et al., 2013; Huang et al., 2015). Thus, we examined whether the dark rearing-induced re-enlargement of RFs could be mediated by a reduction in adult PSD-95 expression. We found that PSD-95 protein levels were not significantly different in DR (0.613± 0.96, n=10) compared to NR adult hamsters (0.486 ± 0.868, n=9) (U(31)=-0.978, p=0.270, Mann-Whitney Rank Sum test) (**Figure 6B**). These results suggest that differences in PSD-95 levels do not underlie the re-enlargement of RFs in SC following dark rearing from birth.

### Cation-chloride co-transporters undergo their normal developmental switch in adult dark reared subjects

Levels of inhibitory GABAergic signaling in neurons are dependent on the intracellular chloride (Cl^-^) concentration. The K^+^ Cl^-^ co-transporter (KCC2) is responsible for regulating intracellular Cl^-^ in mature adult neurons with an outward K^+^ current (Rivera et al., 1999) and also regulates the formation, function, and plasticity of glutamatergic synapses (Li et al., 2007; Gauvain et al., 2011; Chevy et al., 2015). At the beginning of postnatal life, GABA_A_ receptor effects are excitatory because the Na^+^-K^+^-2Cl^−^ co-transporter 1 (NKCC1) that mediates Cl^−^ uptake is dominant (Cherubini et al., 1991; Lee et al., 2005; Cancedda et al., 2007). By the end of the second postnatal week in rats and mice NKCC1 is replaced by KCC2 as the dominant cation-chloride co-transporter in the brain, shifting the resting membrane potential and thus causing GABA_A_ receptors to produce inhibitory PSPs (Rivera et al., 1999; Pfeffer et al., 2009; Moore et al., 2019). In V1, the developmental switch from NKCC1 dominance to KCC2 dominance occurs at the same time as a period of BDNF/TrkB mediated synaptic imbalance (Zhang et al., 2018). We surmised that a shift in the ratio of KCC2:NKCC1 could underlie the reinstatement of RF size plasticity in dark reared adults, leading to re-enlargement of RFs in SC. Examination of the expression of KCC2 and NKCC1 in adult SC neurons revealed no significant differences between our treatment groups, however. KCC2 levels were not significantly different between NR (0.979 ± 0.115, n=8) and DR groups of adult hamsters (0.963 ± 0.154, n=8) (T(14)=0.082, p=0.936, t-test) (**Figure 7A**). The same was true of NKCC1 levels (NR 1.050 ± 0.0419, n=8; DR 1.081 ± 0.0814, n=8) (T(14)=-0.339, p=0.740, t-test) (**Figure 7B**), and of the ratio of the two cation-chloride co-transporters within groups (T(8)=1.096, p=0.305, t-test) (Figure 7C).

**Figure 7.** Cation-chloride co-transporter expression in SC is not affected by early dark rearing. **Images**: Example Western blots of NR and DR samples labeled for cation-chloride co-transporters **(A)** KCC2 (140 kDa) and **(B)** NKCC1 (150 kDa) compared to the GAPDH loading control and **(C)** a comparison of the within subject ratio of KCC2:NKCC1 in NR and DR adult hamsters. **Plots:** Boxplots showing the levels of KCC2 and NKCC1 proteins, normalized against GAPDH **(A and B, respectively)** and comparison of normalized values of KCC2/GAPDH to that of NKCC1/GAPDH **(C)**. Boxes in each boxplot show the median, 25^th^ and 75^th^ percentiles of the data. Whiskers are at 5 and 95% percentiles. Individual data points obtained in each group are shown as dots. These results show that the number and ratio of cation chloride co-transporters are similar in normal and dark reared adult SC.

## Discussion

The goal of this study was to examine potential postsynaptic mechanisms through which light exposure during an early critical period ensures the long-term stability of visual receptive fields in the hamster superior colliculus. Our previous results established a correlation between the maintenance of RF refinement and levels of GABA immunoreactivity in SC (Carrasco et al., 2011; Mudd et al., 2019) and V1 (Balmer and Pallas, 2015), but potential postsynaptic changes in GABA_A_ receptor and related protein levels had not been examined. We have reported here that at the high-resolution level of Western blot protein quantification, visual deprivation-induced failure to maintain refined RFs in SC does not appear to involve changes in GABA_A_R subunit composition, inhibitory or excitatory scaffolding protein expression, or cation-Chloride co-transporter ratios. These results exclude several possible mechanisms that could explain the reduced activation of GABA_A_Rs with GABA agonists reported in DR adult SC (Carrasco et al., 2011), and support activity-dependent regulation of GABA expression as the primary mechanism underlying TrkB-mediated maintenance of RF refinement (Mudd et al., 2019). The finding that a change in GABA levels could affect RF refinement in adulthood has important implications for the treatment of memory impairments or brain injury.

This study supports our previous research that provided substantial evidence of a novel, maladaptive adult plasticity in which visually deprived hamsters refine SC RFs normally but fail to maintain them in adulthood. Our research differs from these previous studies in suggesting that dark reared hamsters lose visual refinement in adulthood and not, as in the case of monkeys, ferrets, and cats, during development (Mower and Christen, 1985; Mower et al., 1986; Mower, 1991; Carmignoto and Vicini, 1992; Fagiolini et al., 1994; Chen et al., 2000; Chen et al., 2001; Lee and Nedivi, 2002; Erisir and Harris, 2003). Some of these previous studies looked only at early and/or adult ages in the animals, thereby missing the RF refinement that happens in between the two ages. Because diminished GABA release, contrary to what we expected, did not elicit measurable changes in the levels of postsynaptic GABA_A_ receptors, scaffold proteins, or chloride co-transporters, this study provides a valuable demonstration that changes in neurotransmitter availability do not necessarily result in coordinated changes in postsynaptic receptors.

Maturation of GABAergic signaling in visual cortex, particularly of the fast-spiking, parvalbumin-containing basket cells, is thought to open and then close the critical period for plasticity (Fagiolini et al., 2004; Sale et al., 2010; Toyoizumi et al., 2013; Capogna et al., 2021). Combined pre- and postsynaptic alteration of synaptic strength has been seen in other sensory deprivation paradigms, including in dark reared and monocularly deprived visual cortex, although with an earlier time course (Carmignoto and Vicini, 1992; Chen et al., 2000; Chen et al., 2001). However, the retinorecipient layers of the superior colliculus have no basket cells and contain very few GABAergic parvalbumin neurons, and the plasticity described here occurs after the critical period has closed, suggesting that SC may accomplish plasticity through a different mechanism than visual cortex. On the other hand, previous studies found that, as in visual cortex (Hanover et al., 1999; Huang et al., 1999; Viegi et al., 2002), deprivation-induced receptive field plasticity in adult SC is mediated by the BDNF receptor TrkB (Mudd et al., 2019). Furthermore, reduced GABA and GABA_A_ receptor efficacy in response to iontophoretically-applied GABA_A_R agonists and antagonists is observed in both SC and V1 of dark reared hamsters (Carrasco et al., 2011; Balmer and Pallas, 2015), arguing for mechanistic elements in common.

### Early visual experience is not necessary for maturation or maintenance of GABA_A_ receptor subunit composition at the synapse

GABA_A_ receptors contain fast acting chloride (Cl^-^) channels (Pfeiffer et al., 1982; Sigel and Steinmann, 2012). The subunit composition of GABA_A_ receptors changes during development from an alpha 2 to alpha 1 dominant condition (Laurie et al., 1992; Fritschy et al., 1994; Chen et al., 2001) and also changes in some disease states (Levitt, 2005; Tyson and Anderson, 2014; Deidda et al., 2015; Kimoto et al., 2015; Schmidt and Mirnics, 2015; Tang et al., 2021) in a way that affects receptor functional properties (Farrant and Nusser, 2005) and localization (Jacob et al., 2005). We studied synaptic levels of the GABA_A_ receptor alpha1 and alpha2 subunits to quantify their expression levels under normal and DR conditions. The normalized expression levels of GABA_A_R alpha1 relative to GABA_A_R alpha2 levels were not altered in DR hamsters when compared to those of NR hamsters, arguing that the altered inhibitory synaptic efficacy that we previously observed was not caused by an immature GABA_A_R subunit composition at the synapse.

An increase or decrease in the level of any receptor subunit is best understood in context, because different conclusions would be drawn if subunits changed independently or in concert. Thus, we also analyzed the alpha1/alpha2 ratios in individual animals. We did not find any change in alpha1/alpha2 ratios in NR compared to DR adult hamsters. These results suggest that early visual experience is not necessary for maturation or maintenance of mature synaptic GABA_A_R subunit composition in adulthood. Thus, the failure to maintain refined RFs in adult DR hamsters cannot be explained by a return to a juvenile type of GABA_A_R subunit composition.

### Level and localization of the extrasynaptic GABA_A_R subunit alpha 5 does not change with dark rearing

GABA_A_R subunit alpha 5 is predominantly an extrasynaptic membrane receptor subunit that regulates tonic inhibition. It is important in neuronal circuit development, learning, and memory (Brady and Jacob, 2015), has been implicated in regulating the induction of synaptic plasticity in hippocampus (Saab et al., 2010; Zurek et al., 2012; Zurek et al., 2014), and can relocate to the synaptic region in learning and memory deficits (Brady and Jacob, 2015). Because the excitation/inhibition (E/I) balance could be affected if alpha5 subunit levels changed or if they moved into the synapse, we compared its expression between NR and DR cases. We did not see any significant changes in the overall levels of GABA_A_R alpha5 subunits, or in the alpha1/alpha5 ratio, suggesting that dark rearing-induced RF enlargement is not caused by changes in the GABA_A_R subunit alpha 5 levels or localization in the synapse.

### Dark rearing does not affect the rate of internalization of GABA_A_R subunits at the synapse

Because clathrin-dependent endocytosis is likely important for regulating inhibitory signaling and synaptic plasticity (Kittler et al., 2000), we explored the internalization of GABA_A_ receptor alpha1 and alpha5 subunits by comparing their synaptic vs. extrasynaptic location in normally reared and dark reared subjects. We did not observe a significant change in location of either subunit type as assayed by the ratio of synaptosome-bound to cytosolic fractions. This implies that a lack of visual experience does not affect the trafficking of the GABA_A_R subunits between the synaptic membrane and the cytosol or the phosphorylation events that maintain the balance between internalization and postsynaptic membrane insertion of the receptor subunits.

### Early visual experience is not necessary to maintain scaffolding protein levels at the synapse

Another finding of this work is that the expression levels of the postsynaptic scaffold proteins PSD-95 and gephyrin were not altered in adulthood following lifelong lack of light exposure, suggesting that any changes in inhibitory function are probably not caused by a significant change in scaffolding protein expression. At any rate, the clustering of GABA_A_Rs at inhibitory synapses in SC may happen in a gephyrin-independent manner (Kneussel et al., 2001), or total gephyrin expression may not be as important as the formation of gephyrin nanodomains within inhibitory synapses (Pennacchietti et al., 2017). Future studies with additional techniques would be required to determine if changes in receptor clustering may be occurring and what role gephyrin or PSD-95 may have in mediating any such effects.

### Early light exposure is not necessary for maturation of the cation-chloride co-transporters

We investigated the status of the chloride transporters KCC2 and NKCC1 due to their role in maintaining chloride balance inside of the neurons and thus in setting the reversal potential. The cation-chloride co-transporters could have reverted to their early developmental state, leading to a lower threshold for excitation in dark reared animals, possibly explaining the RF expansion we observed. However, we did not observe any changes in the cation-chloride co-transporters in dark-reared compared to normally reared adult hamsters, suggesting that the RF enlargement was not caused by alterations in the cation-chloride co-transporters.

### A GABA-BDNF feedback loop maintains inhibitory networks, thereby maintaining RF refinement in adulthood

GABA-GABA_A_R interaction is known to regulate various downstream signaling pathways, and a major regulator of GABA itself is BDNF-TrkB signaling triggered by NMDA receptor activity (Marini et al., 1998). Our data are consistent with previous studies suggesting a positive feedback loop between the BDNF-TrkB pathway and GABA expression, in which GABA facilitates BDNF expression, and BDNF facilitates the production of GABA by GAD (Sánchez-Huertas and Rico, 2010) and its synaptic release (Huang et al., 1999; Morales et al., 2002; Gianfranceschi et al., 2003; Jovanovic et al., 2004; Kuczewski et al., 2008; Porcher et al., 2011; Hanno-Iijima et al., 2015), maintaining RF size and visual acuity through GABAergic lateral inhibition (Mudd et al., 2019). Signaling via the MAPK cascade and the transcription factor cAMP-responsive element-binding protein (CREB) appears to play a substantial role in this process (Obrietan et al., 2002; Sánchez-Huertas and Rico, 2010). BDNF-TrkB interaction leads to dimerization and auto-phosphorylation of TrkB, thereby triggering MAPK, PLC gamma, and PI3K pathways (Yoshii and Constantine-Paton, 2007). These pathways in turn lead to the activation of downstream effectors and mediators to initiate a CREB-dependent transcription process that can lead to an increase in GABA_A_R levels as well as more BDNF production (Huang and Reichardt, 2003; Yoshii and Constantine-Paton, 2007; Porcher et al., 2011; Esvald et al., 2020). In addition, an increase in the transmembrane localization of GABA_A_Rs is mediated by BDNF-dependent inhibition of receptor internalization in addition to ongoing reinsertion of the receptor into the postsynaptic membrane (Porcher et al., 2011). This positive feedback regulation is critical in developing neurons and hence constituted a major motivation for the work reported here. In this study however, neither the GABA_A_R subunit composition at the synapse nor subunit composition in the extrasynaptic regions was affected by dark rearing. Chloride transport proteins (KCC2, NKCC2) also remained at normal levels. One possibility is that GABA expression levels alone are the key factor in RF re-enlargement in hamster SC (Carrasco et al., 2011; Mudd et al., 2019). If so, it would suggest that this type of delayed plasticity resulting from a lack of early visual experience occurs through a different mechanism than described in other types of plasticity resulting from dark rearing (Mower et al., 1985; Chen et al., 2000; Chen et al., 2001).

It is possible that changes in GABA_A_R signaling occurred that are not reflected here in the expression levels of postsynaptic receptor composition, scaffolding molecules, or ion transporters, or that we missed some transient changes in GABA_A_R signaling-associated proteins that cause GABA_A_R functional changes. In the future, it would be interesting to study protein localization and interactions with immunohistochemistry, or to study the properties of synaptic and extrasynaptic responses in the SC with patch clamp experiments, and the subcellular dynamics of associated proteins involved. This might improve the understanding of the molecular processes active in this deprivation-induced, maladaptive plasticity in the SC.

Suggested alternative explanations include that the changes leading to RF expansion and thus the visual acuity deficits take place earlier than the time point that we studied and return to normal by P90. The width of the synaptic cleft decreases during development (Li and Cline, 2010), thus, another interesting possibility is that dark rearing from birth gradually increases the width of the synaptic cleft in adulthood while keeping the postsynaptic signaling components in place. Alternatively, the synapses may be present but silent due to the presynaptic loss of GABA (Carrasco et al., 2011). It is also possible that other scaffolding proteins and their partners are involved. Potential candidates include gephyrin binding partners such as GABA_A_R beta2 and beta3 subunits (Kowalczyk et al., 2013), the scaffold protein radixin that binds GABA_A_R alpha5 subunit to the actin cytoskeleton (Loebrich et al., 2006) and regulates synaptic GABA_A_R density (Hausrat et al., 2015), or gephyrin post-translational modification events that influence inhibitory synaptic plasticity by affecting postsynaptic scaffolding (Zacchi et al., 2014). More recently, distinct spectrin isoforms have been shown to affect synaptic inhibition by selectively targeting specific GABA_A_R subunits, including α1 and α2, to particular regions of the neuron (Smalley et al., 2023).

In summary, our results argue that visual experience is not necessary to maintain mature levels and composition of several postsynaptic proteins that are essential for retino-SC synaptic communication. Unlike many previous dark rearing studies in which both GABA and GABA_A_ receptors were found to be downregulated (Carmignoto and Vicini, 1992; Chen et al., 2000; Chen et al., 2001; Kilman et al., 2002; Nahmani and Turrigiano, 2014), we report here that in the SC, GABA_A_R levels, subunit composition, and localization in adulthood are unaffected by dark rearing. The scaffold proteins gephyrin and PSD-95, and the chloride transporters KCC2 and NKCC2 are also not affected by dark rearing. This novel, experience-dependent form of adult plasticity may involve an as yet unidentified postsynaptic mechanism, or only a reduction in GABA release (Carrasco et al., 2005; Carrasco and Pallas, 2006; Carrasco et al., 2011), thereby challenging the common view that presynaptic changes in ligand availability are always associated with matching postsynaptic changes in their receptors. Either possibility is encouraging with respect to understanding this form of adult plasticity and might help adults with memory impairments, traumatic brain injury, or inhibition-associated neurological disorders.

## Acknowledgments

We thank Profs. Angela Mabb, Larry Schwartz, Thomas Maresca, and Lillian Fritz-Laylin for instrument access and guidance with Western Blotting, Profs. Gerald Downes, Margaret Stratton, and Stephen Moss for providing technical guidance and support on P.S.J.’s thesis committee, Pallas lab members for technical support and manuscript review, and the animal care staff at GSU and UMass. We thank Dr. Andrew Stephens for providing the Histone H3 antibody.

## Conflict of interest

*The authors declare that the research was conducted in the absence of any commercial or financial relationships that could be construed as a potential conflict of interest*.

## Author Contributions

Juvale, Mudd, and Pallas contributed to the conception and design of the study. Juvale, Mudd, and Shree collected data, organized the database, and performed the statistical analyses. Mudd wrote the first draft of the manuscript. Juvale wrote subsequent versions of the manuscript and Pallas edited the manuscript. All authors read the manuscript, contributed to manuscript revision, and approved the submitted version.

## Funding

Support for this work was provided by a Sigma Xi GIAR grant and a UMass Amherst predissertation grant to P.S.J.; a GSU Brains & Behavior Fellowship, and a GSU Center for Neuromics student grant awarded to D.B.M.; and a GSU Brains & Behavior Seed grant, UMass startup funds, a National Science Foundation grant (IOS-1656838), and a DARPA grant (HR0011-18-2-0019, TA2) awarded to S.L.P.

## Notes

### Competing Interest Statement

The authors have declared no competing interest.

### Summary of Updates

Minor changes in the text, ORCID IDs of authors, addition of references.

